# Diversity of Lepidoptera in urban parks at Pasir Gudang, Johor, Malaysia

**DOI:** 10.1101/2021.11.29.470299

**Authors:** Sharifah Fatin Naqliah Syed Nisranshah, Nurulhuda Zakaria

## Abstract

Urban parks are a valuable habitat for Lepidoptera, which are often neglected despite their potential role as pollinators and important food source for small animals. Lepidoptera in urban parks in Pasir Gudang are still under-studied. Yet, there is no publication on Lepidoptera in these urban parks. The objectives of this study were to determine and to compare the diversity of Lepidoptera in three urban parks (Laman Rekreasi Sungai Buluh, Tasik Dahlia and Taman Bandar) at Pasir Gudang, Johor. Visual Encounter Survey (VES) method was used to capture the Lepidoptera from February 2021 until April 2021. Four sampling sessions were conducted for each study site. A total of nine species from 117 individuals comprised of family Pieridae (43%), Nymphalidae (29%) and Lycaenidae (28%) were recorded from this study. The most abundant species were *Zizula hylax* and *Elymnias hypermnestra*, while *Leptosia nina* was the least abundant. The results from this study showed that the diversity of Lepidoptera have established in these three urban parks. Diversity of Lepidoptera in Tasik Dahlia was significantly lower compared to the other two study sites. The findings from this study could be useful as a baseline data for future research and conservation management of the order Lepidoptera around the urban parks and indirectly can support the sustainable development of the urban parks.

## INTRODUCTION

One of the most commonly known as beautiful creature in the world is butterfly. The beautifulness of butterfly adds more favour in scenery and such an eye candy. What gives the butterfly its elegance are the scales, which are arranged in colourful designs unique to each species. The order Lepidoptera belongs to moths and butterflies. Lepidoptera is locally known as “rama-rama” and “kupu-kupu” in Malaysia. Globally, more than 150,000 species have been described from this order (Gullan et al., 2010). Among the main characteristics of the Lepidoptera are high mobility, quick perception of stimuli and has a strong exoskeleton. There are distinctive morphology and biological behaviours between moths and butterflies even they are in the same order in taxanomy hierarchy.

Lepidoptera have role to ensure the food webs are continuously run on the right path in the ecosystem they live. Lepidoptera are important pollinators for many plants (Andersson et al., 2002). Lepidoptera can serve as food source for insectivorous vertebrates such as birds (Gullan et al., 2010). Lepidoptera can be considered as keystone species because any loss of their critical ecological functions could collapse the entire ecosystem. Lepidoptera are suitable to be chosen as the subject for this study as they are a very good indicators of the health of the ecosystem (Dennis et al., 2017).

Lepidoptera are very sensitive toward their environment even with the small changes surrounding them. The biggest threat to the order Lepidoptera are residential and commercial developments which involve housing & urban areas, tourism and recreation areas that can contribute to pollution and habitat shifting (IUCN Redlist, 2021). Moreover, industrialization and urbanization also changed the natural habitat or niches for the plants and animals. Pasir Gudang is a city in the district of Johor Bahru, Johor, Malaysia. The major industries in this city are transportation and logistics, shipbuilding, petrochemicals and other heavy industries. The parks within this city are considered as urban parks which are the places that offer recreation and green space to the residents. The major concern here is whether the Lepidoptera are able to survive in this contaminated urbanized area.

These urban parks continuously received various degree of environmental stress such as air pollution that come from numerous factories that release toxic gases to the air or exhaust gases from transportation, noise pollution or light pollution. The survival of Lepidoptera in contaminated urbanized area is still questionable until this day. All of these pollutions might disturb moth and butterflies behaviours and preferences such as feeding, mating and life cycle. According to Yap et al. (2019), Pasir Gudang is one of the hotspots of contaminated area in Johor, Malaysia. This might give big impact and lead to the modification of characteristics or new traits to the organisms so that they can adapt with anthropogenic disturbance to ensure the continuous survival of the next generations.

The major aims of this study were to determine and compare the diversity of Lepidoptera between three urban parks in Pasir Gudang, Johor. Taman Bandar, Laman Rekreasi Sungai Buluh and Tasik Dahlia are still under-studied urban parks in Pasir Gudang, Johor Malaysia. Yet, there is no publication on species composition in these urban parks. In this study, the composition of species was recorded to give an early idea on how the urban parks have influenced the pattern of the Lepidoptera’s diversity. At the same time, the species richness also was determined. The identification of Lepidoptera in this study will contribute to the first database of Lepidoptera in the three urban parks in Pasir Gudang, Johor and assessment of their biodiversity and conservation status can be made. This study will also help people to gain more knowledge about the order Lepidoptera around the urban areas and may give important information about ecological roles of Lepidoptera.

## Materials and Methods

### Study sites

This study was carried out at Pasir Gudang, Johor that is located in the southern region of the Peninsular Malaysia (1°28’13.1”N, 103°54’10.8”E) (Figure 1). The selected sampling sites are located in Pasir Gudang town that is located 26.3 km away from Johor Bahru city. Tasik Dahlia, Laman Rekreasi Sungai Buluh and Taman Bandar are the three urban parks in Pasir Gudang town that were selected as sampling sites. The detail descriptions about each of the urban park are shown in Table 1. These urban parks are surrounded primarily by residentials, restaurants, main roads, grocery-stores and buildings. These urban parks play an important role to public community in ensuring the public keep in touch with the nature and have a balance and healthy lifestyle.

**Table 1.**
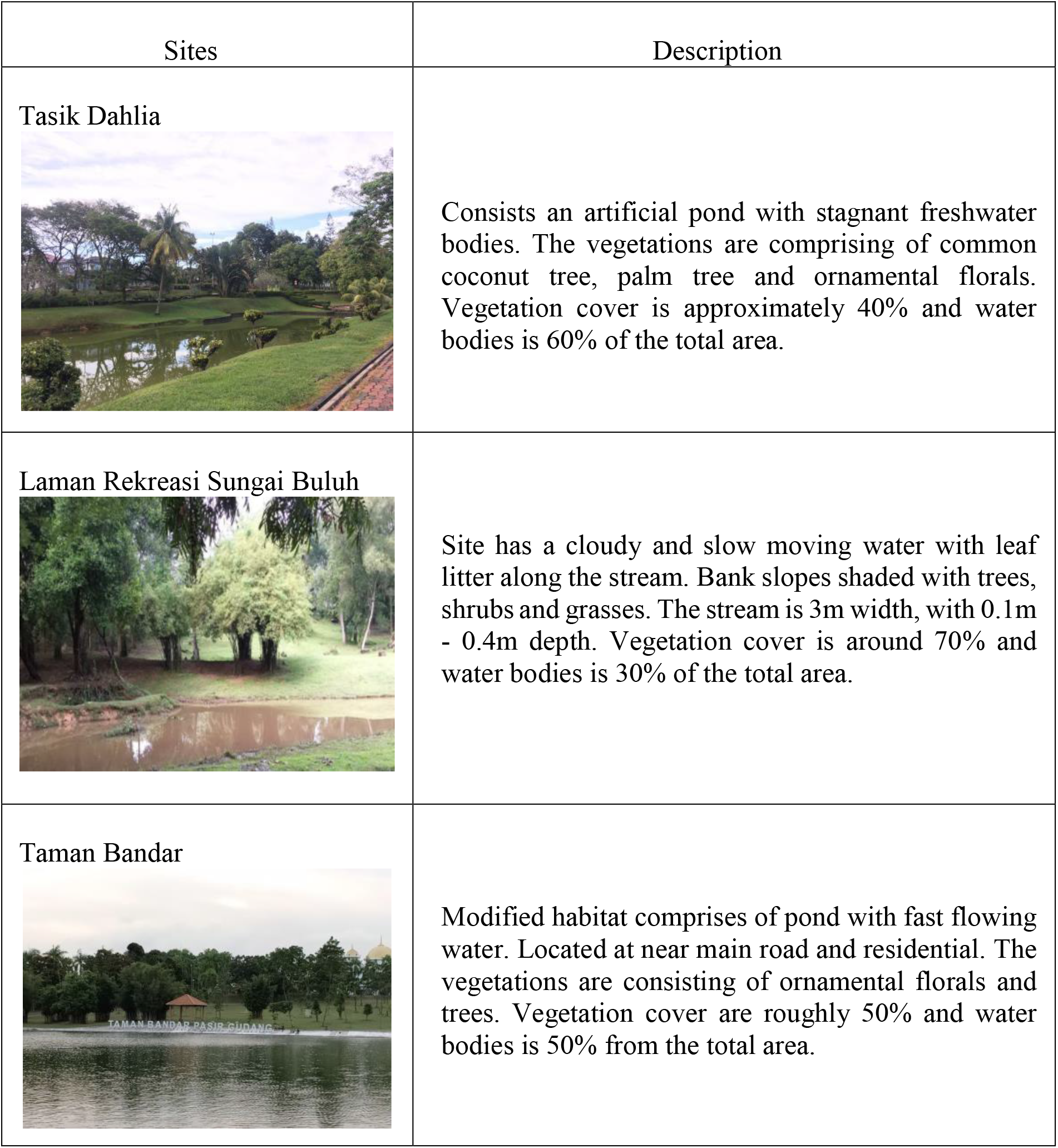
The description of the physical characters of the study sites.

| Sites | Description |
| --- | --- |
| Tasik Dahlia | Consists an artificial pond with stagnant freshwater bodies. The vegetations are comprising of common coconut tree, palm tree and ornamental florals. Vegetation cover is approximately 40% and water bodies is 60% of the total area. |
| Laman Rekreasi Sungai Buluh | Site has a cloudy and slow moving water with leaf litter along the stream. Bank slopes shaded with trees, shrubs and grasses. The stream is 3m width, with 0.1m - 0.4m depth. Vegetation cover is around 70% and water bodies is 30% of the total area. |
| Taman Bandar | Modified habitat comprises of pond with fast flowing water. Located at near main road and residential. The vegetations are consisting of ornamental florals and trees. Vegetation cover are roughly 50% and water bodies is 50% from the total area. |

**Figure 1.**
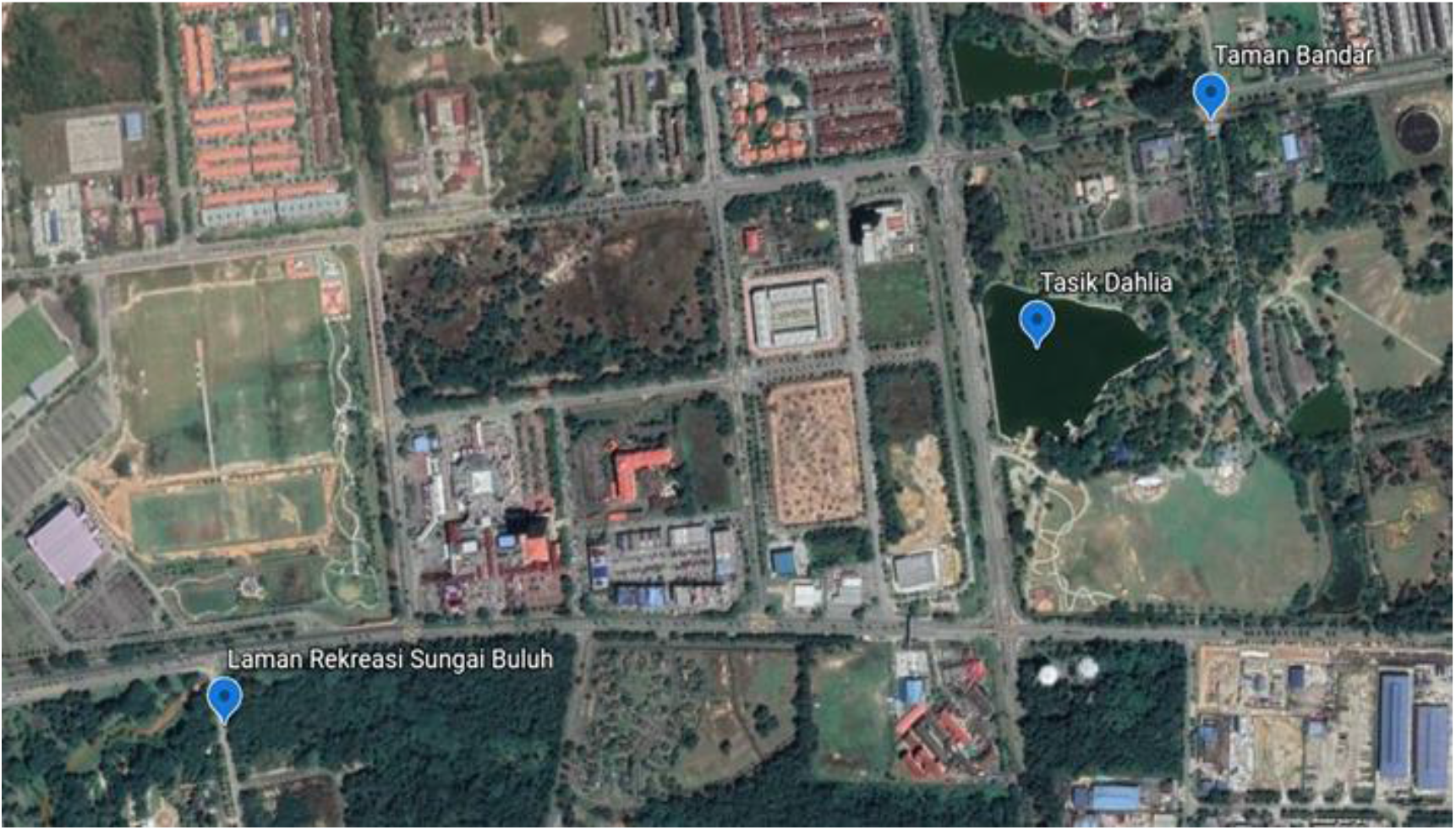
The location of three sampling sites were shown in blue dots (Google Maps, 2020).

### Methods

In this study, Visual Encounter Survey (VES) was chosen as the sampling method. Visual Encounter Survey is an active sampling, by definition a time-constrained method in which observers sample for species richness and abundance along a survey path (Crump & Scott, 1994). The VES was carried out during specific time which was six hours (0800 am – 1100 am and 0200 pm – 0500 pm) for each site. Data collection for this project was performed on monthly basis from February 2021 until April 2021. Sampling session for each site was conducted four times per sampling site. Overall, a total of 12 times sampling sessions were executed within three months. Samplings were carried out on the existing trail at the urban parks. Lepidoptera were captured using a sweep net. Fruits were used as baits to attract the Lepidoptera. Each of the captured animals were placed in a zip lock bag, identified, photographed and released back into the wild. Animals were identified using Kirton (2020) and iNaturalist (2021).

### Data analysis

The ecological indices of diversity, richness and evenness were calculated for Lepidoptera community in urban parks at Pasir Gudang, Johor. The Shannon Index and Simpson Index are the diversity metrics that were used to determine the diversity of Lepidoptera in this study. Clarke and Warwick (2001) stated that the Shannon-Weiner Index is the common index widely used to compare species diversity between different habitats. The Simpson index is the dominance index because it gives the common or the dominant species more weight. Margalef and Menhinick indices were measured to estimate the species richness. Species richness is the number of species present in a region that can be counted simply as the number of different species. It does not take into account the species abundance or its relative distribution of abundance. The Margalef scale is the simplest way to measure resources of biodiversity (Margalef, 1958). The Menhinick index was calculated using a formula from Mirzae et al. (2013). Species evenness was determined by using Simpson Index. Shannon Index (H’) was used to calculate the species diversity based on the number of individuals and number of species.

Rarefaction is recognized from the results of the sampling as a tool for determining species resources. Rarefaction is a method for estimating the species richness in a small sample when abundance information is given for a larger sample (Krebs, 1989). This approach can be used to compare the number of species in different samples. In this study, the number of species and individuals were used to construct the rarefaction curve. Meanwhile Rank abundance curve has been used to display the relative abundance of the species. In this study, the rank abundance curve was constructed using the number of individuals of a species and the number of species. The accumulation curve of species is used to measure or estimate the number of species in a given area (Hammer et al., 2001). Species accumulation curve allows to evaluate number of species that could be found in a given area and to compare the diversity across population or to determine the benefits of additional sampling. The sampling will be considered as maximum sampling when the species accumulation curve is levelling off which indicates no sampling effort are needed. In this study, the species accumulation curve was constructed using the number of species and number of individuals.

All diversity indices were calculated by using Paleontological Statistic (PAST) software (Hammer et al., 2001) while Rarefaction curve, Rank abundance curve, Species accumulation curve were constructed using EcoSim700 software (Gotelli & Entsminger, 2004).

### Statistical analysis

Diversity t-test was performed in this study by using Paleontological Statistic (PAST) software (Hammer et al., 2001). Diversity t-test was used to detect the significant differences in species diversity among three different sites. A significant different means that the result that are most likely to be seen are not due to a random or sampling error. Diversity t-test was used to compare the diversity of Lepidoptera between three different urban parks in Pasir Gudang city through a pairwise comparison. Diversity t-test was conducted using Shannon-Weiner and Simpson indices values. The p-value < 0.05 indicated that there was a significant different in the diversity of Lepidoptera between the three urban parks.

## RESULT

### Composition of Lepidoptera

A total of 117 Lepidoptera individuals were captured, comprising of three families and nine species as listed in the Table 2. The composition of Lepidoptera according to the family were Lycaenidae (three species), Nymphalidae (two species) and Pieridae (four species). The most abundant family was Pieridae with four species recorded while the least abundant family was Nymphalidae with only two species. The highest species number was recorded at Laman Rekreasi Sungai Buluh urban park with a total of nine species and the lowest was at Tasik Dahlia urban park with only four species. The highest number of individuals was recorded from the Laman Rekreasi Sungai Buluh (117 individuals) while the least was at Tasik Dahlia (28 individuals).

**Table 2.**
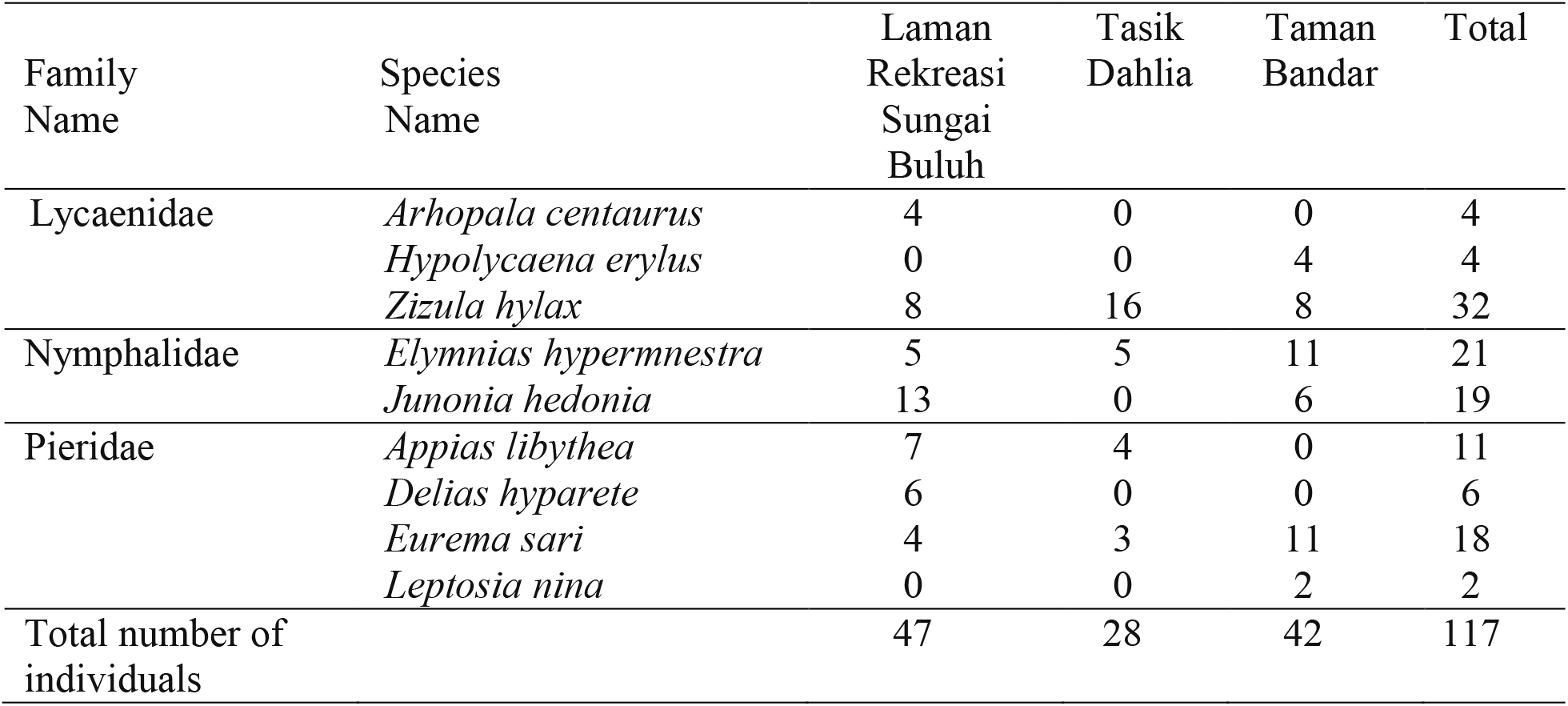
Composition of Lepidoptera at three different urban parks at Pasir Gudang, Johor.

| Family Name | Species Name | Laman Rekreasi Sungai Buluh | Tasik Dahlia | Taman Bandar | Total |
| --- | --- | --- | --- | --- | --- |
| Lycaenidae | *Arhopala centaurus* | 4 | 0 | 0 | 4 |
|  | *Hypolycaena erylus* | 0 | 0 | 4 | 4 |
|  | *Zizula hylax* | 8 | 16 | 8 | 32 |
| Nymphalidae | *Elymnias hypermnestra* | 5 | 5 | 11 | 21 |
|  | *Junonia hedonia* | 13 | 0 | 6 | 19 |
| Pieridae | *Appias libythea* | 7 | 4 | 0 | 11 |
|  | *Delias hyparete* | 6 | 0 | 0 | 6 |
|  | *Eurema sari* | 4 | 3 | 11 | 18 |
|  | *Leptosia nina* | 0 | 0 | 2 | 2 |
| Total number of individuals |  | 47 | 28 | 42 | 117 |

### Rank abundance Curve

Based on Figure 2, the most abundant species at Tasik Dahlia was *Zizula hylax* with 16 individuals. The most abundant species recorded at Laman Rekreasi Sungai Buluh was *Junonia hedonia* with 13 individuals. Tasik Dahlia showed the steepest slope thus indicated that it had the lowest evenness compared to the Laman Rekreasi Sungai Buluh and Taman Bandar.

**Figure 2.**
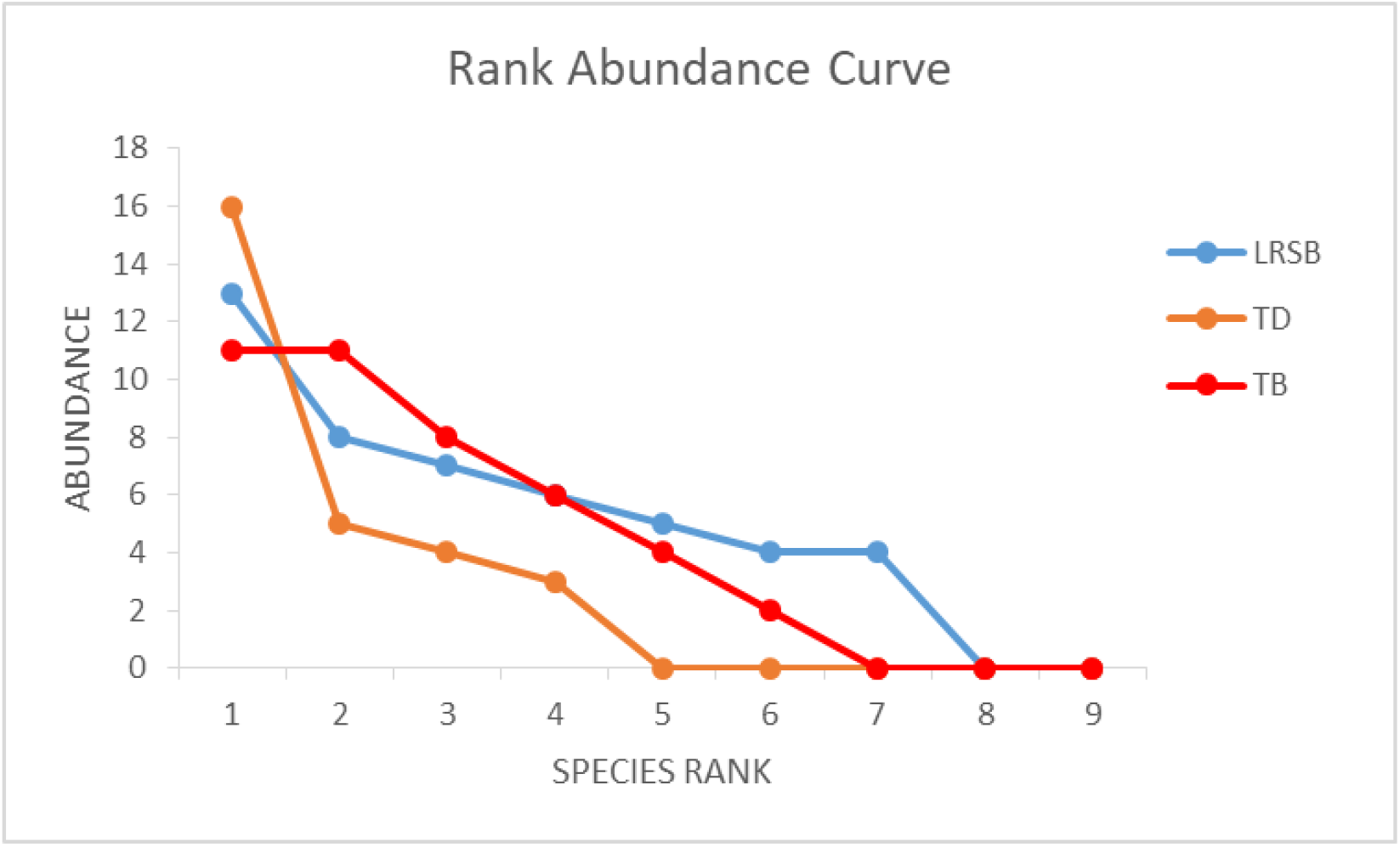
Rank abundance curve of each urban park at Pasir Gudang, Johor. Note: LRSB= Laman Rekreasi Sungai Buluh, TB= Taman Bandar, TD= Tasik Dahli

### Species accumulation curve

Figure 3 showed the species accumulation curves at three different urban parks in Pasir Gudang, Johor. Curves for all three urban parks had reached a plateau indicating that the sampling efforts were enough, and no further species will be recorded although more samplings sessions are added. However, the highest species accumulation curve from Laman Rekreasi Sungai Buluh indicated an inadequate number of individuals were captured and this showed that this urban park had higher number of species been caught compared to Taman Bandar and Tasik Dahlia.

**Figure 3.**
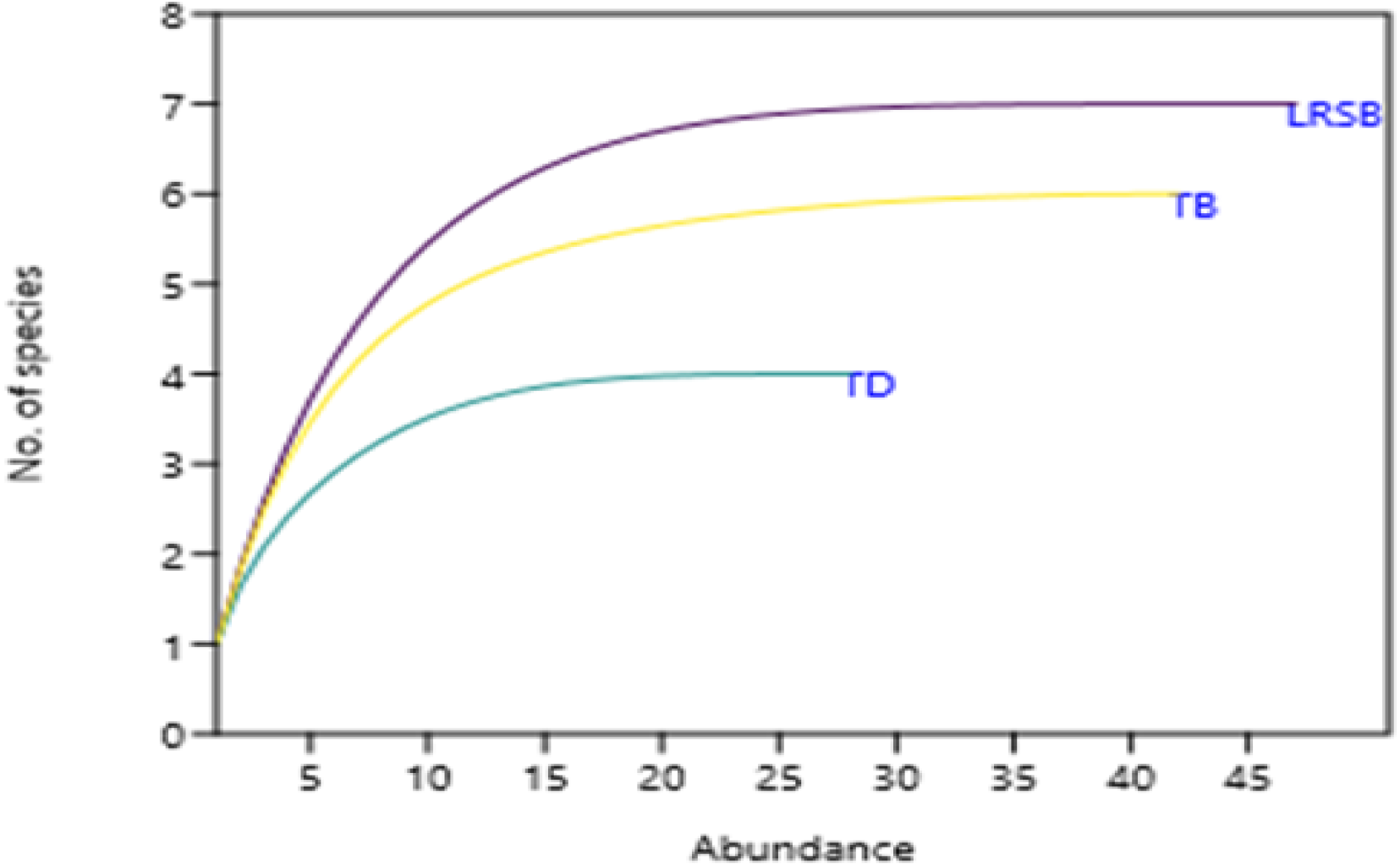
Species accumulation curves for each urban park at Pasir Gudang, Johor. Note: LRSB= Laman Rekreasi Sungai Buluh, TB= Taman Bandar, TD= Tasik Dahlia

### Rarefaction curve

Figure 4 showed an individual based rarefaction curve to compare the diversity of species captured at three urban parks at Pasir Gudang, Johor. Rarefaction curves for all three sites grew rapidly at first, as the most common species were found and then the curves became plateau as only the rarest species remain to be sampled.

**Figure 4.**
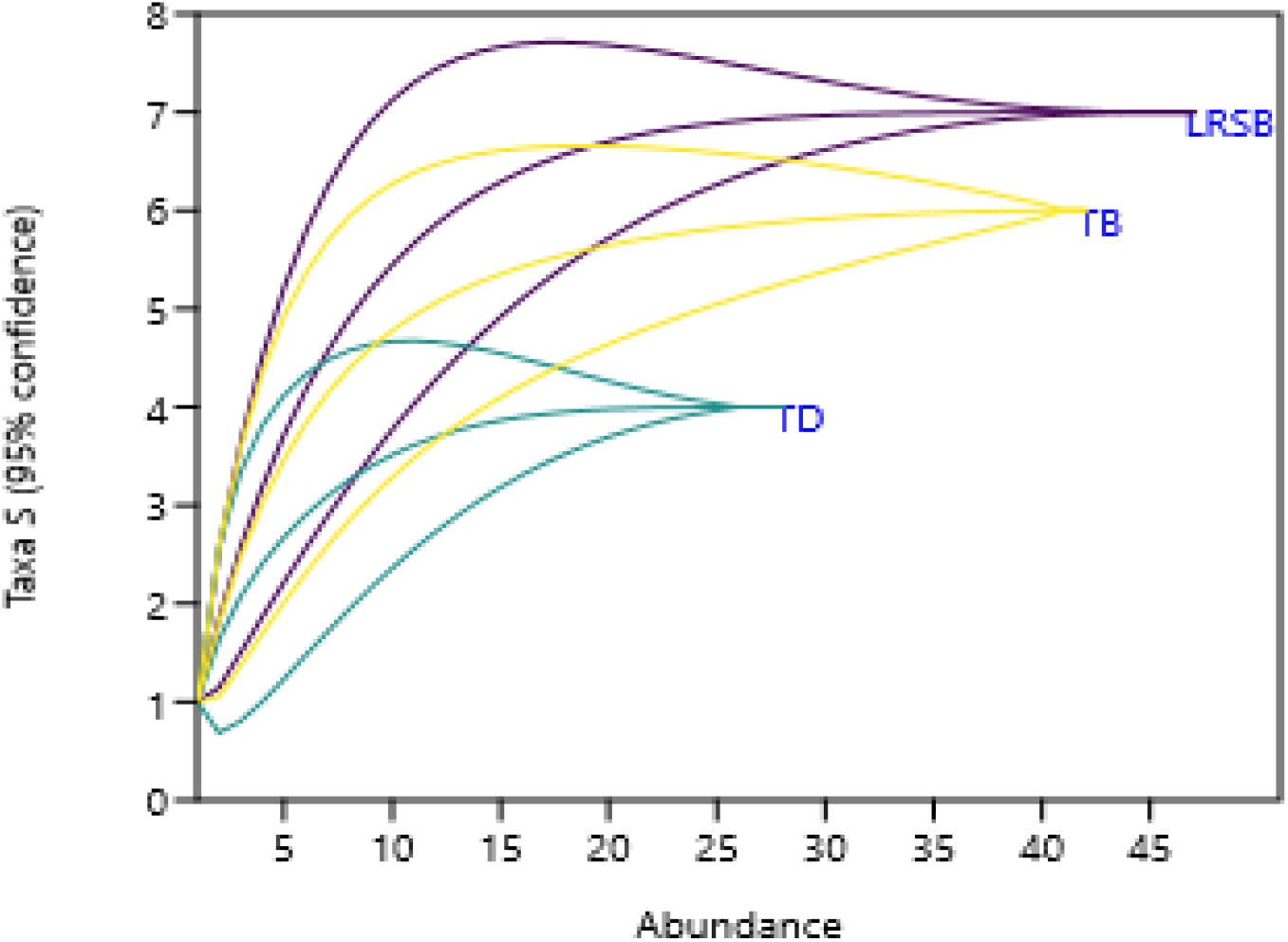
Rarefaction curves for each urban park at Pasir Gudang, Johor. Note: LRSB= Laman Rekreasi Sungai Buluh, TB= Taman Bandar, TD= Tasik Dahlia

### Diversity indices

Based on Table 3, the highest number of species was recorded at Laman Rekreasi Sungai Buluh with seven species from 47 individuals. Tasik Dahlia recorded the least with only four species with 28 individuals. This implied that Tasik Dahlia sampled the lowest species evenness and lowest diversity of Lepidoptera species. Tasik Dahlia recorded the highest dominance value (0.3903) compared to the other two study sites.

**Table 3.**
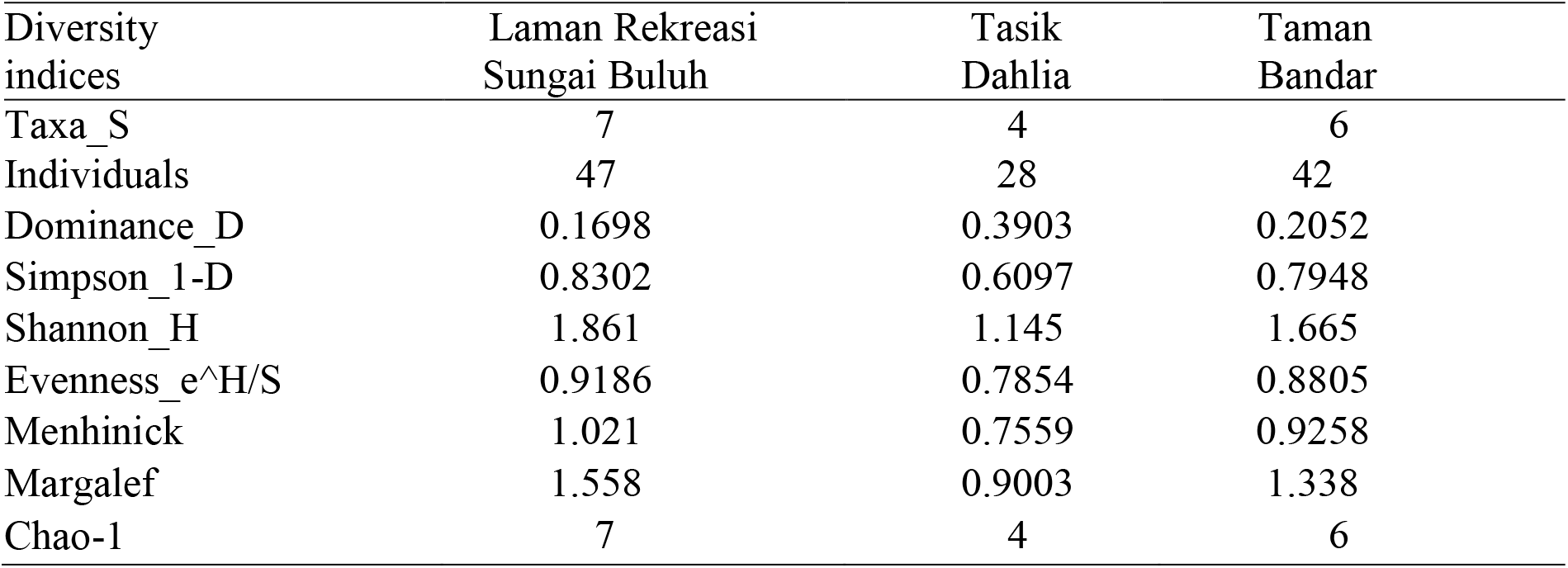
Diversity indices of Lepidoptera in three urban parks at Pasir Gudang, Johor.

| Diversity indices | Laman Rekreasi Sungai Buluh | Tasik Dahlia | Taman Bandar |
| --- | --- | --- | --- |
| Taxa_S | 7 | 4 | 6 |
| Individuals | 47 | 28 | 42 |
| Dominance_D | 0.1698 | 0.3903 | 0.2052 |
| Simpson_1-D | 0.8302 | 0.6097 | 0.7948 |
| Shannon_H | 1.861 | 1.145 | 1.665 |
| Evenness_e^H/S | 0.9186 | 0.7854 | 0.8805 |
| Menhinick | 1.021 | 0.7559 | 0.9258 |
| Margalef | 1.558 | 0.9003 | 1.338 |
| Chao-1 | 7 | 4 | 6 |

For the diversity, Simpson Index was recorded the highest at Laman Rekreasi Sungai Buluh (0.8302) while the lowest was recorded at Tasik Dahlia (0.6097). This trend was also similar with the Shannon index. According to Magurran et al. (2004), Shannon Index which had value of higher than 3.5 showed a high species diversity, value of 1.5 to 3.5 represented moderately high diversity and value of below than 1.5 indicated low in diversity. Between these three study sites, Laman Rekreasi Sungai Buluh and Taman Bandar showed high value of Shannon Weiner index which were 1.861 and 1.665, respectively which indicated moderately high diversity compared to Tasik Dahlia with value of 1.145 which indicated low in diversity.

The highest evenness index value was recorded at Laman Rekreasi Sungai Buluh with 0.9186. Menhinick and Margalef indices shared the similar trend as both represented the highest value among all the study sites. Chao-1 estimated the number of species that can be collected at each of the urban park. When comparing the data from the Table 2 and Table 3, there was no difference between the number of species collected from the samplings with the Chao-1 estimation with Laman Rekreasi Sungai Buluh collected seven species, Tasik Dahlia collected four species and Taman Bandar collected six species. This finding supported the species accumulation curve (Figure 3) that had estimated the sampling efforts were enough and no need to be added in the future in order to get more species of Lepidoptera at all three study sites.

### Comparison of diversity

Diversity of Lepidoptera at Tasik Dahlia showed a significant different with the other two study sites as the *p*-value was below than 0.05 (Table 4). Besides, Laman Rekreasi Sungai Buluh and Taman Bandar recorded no significant different with the two study sites as the *p*-value was above 0.05.

**Table 4.** The *p*-value from the diversity t-test using Shannon Index (upper corner) and Simpson Index (lower corner).

| Study sites | Laman Rekreasi Sungai Buluh | Tasik Dahlia | Taman Bandar |
| --- | --- | --- | --- |
| Laman Rekreasi Sungai Buluh |  | 0.000033291 (S) | 0.068261 (N.S) |
| Tasik Dahlia | 0.014211 (S) |  | 0.0019623 (S) |
| Taman Bandar | 0.30336 (N.S) | 0.037718 (S) |  |

Note: Significant (S) *p*-value < 0.05, not significant (N.S) *p*-value > 0.05

## DISCUSSION

The most abundant species at Laman Rekreasi Sungai Buluh was *Junonia hedonia* from the family Nymphalidae. These could be due to the fact that vegetation structures at Laman Rekreasi Sungai Buluh were numerous with shaded trees, shrubs and grasses. Family Nymphalidae feeding preferences varies between species, with some preferring flower nectar while others preferring sap flows, rotting fruit, dung, or animal carcasses (MyBIS, 2021). The most abundant species at Tasik Dahlia was *Zizula hylax* from the family of Lycaenidae. Overall, the landscape of Tasik Dahlia predominantly covered by open grassy area. *Zizula hylax* is commonly interact with perennial plants. The example of perennial plants species that have interactions with *Zizula hylax* are *Tridax procumbens* and *Cyanthillium cinereum* which can be found at open grassy area (The Biodiversity of Singapore, 2021). The most abundant species at Taman Bandar were *Elymnias hypermnestra* from the family Nymphalidae and *Eurema sari* from the family Pieridae. *Elymnias hypermnestra* used *Cocos nucifera* as a host plant which the plant can be found at Taman Bandar site (MyBIS, 2021). Taman Bandar site is also comprising with fast flowing water pond that could attracted the *Elymnias hypermnestra*. The vegetations at Taman Bandar are consisting of ornamental floral and trees which provide shaded areas. *Elymnias hypermnestra* were often be seen flying along the border of a vegetated area to visit shaded flowering plant in order to get minerals and energy. The specific species of plant that were mostly used as perching spot by *Elymnias hypermnestra* was *Caryota mitis* which can be found at Taman Bandar site. Thus, species richness and species diversity indirectly increased with the increasing of vegetative structures. For *Eurema sari* from the family Pieridae, this species presented mostly at water bodies at Taman Bandar site. *Eurema sari* is commonly be seen mud-pooling or habit of puddling meaning here visitation of mud puddles and patches of moist soil (The Biodiversity of Singapore, 2021).

Figure 4 showed the rarefaction curves at three study sites which were Laman Rekreasi Sungai Buluh, Tasik Dahlia and Taman Bandar. The curve for all three study sites grew rapidly at first as the most common species were found, but then the curves became plateau as only the rarest species remain to be sampled. It can display precipitous curve if or when the evenness of the relative abundance distribution is elevated (Gotelli & Graves, 1996). Rarefaction with the non-overlapped 95% confident interval showed that there were significant differences in the diversity of Lepidoptera between the three study sites. This showed that when the curve reached asymptote, it represented that all existing species in that area were already found. All three curves levelled to a plateau, indicating complete sampling for all three study sites as it did not overlap with each other. According to Aqilah et al. (2019), there were variation of genus and species from the family of Nymphalidae, Lycaenidae and Pieridae that were recorded from the forest reserves in Taka Melor Amenity Forest and Labis Forest Reserve, Segamat, Johor. The checklist from their study showed that the species from these three families have the capability to survive in multifariousness environment and ecosystem especially in disturbed and undisturbed area. Lepidoptera are extremely sensitive to changes in its environment, even if it is a minor change. Residential and commercial development, which involves housing and urban areas, tourism and recreation areas that can contribute to pollution and habitat shifting, pose a major threat to the order Lepidoptera. The natural habitats or niches for plants and animals have changed as a result of industrialization and urbanization. Based on Table 2, family Pieridae had the highest composition of species according to the family at the three urban parks in Pasir Gudang city. This might be influenced by the ability of this family in mimicry inedible species that serve as warning sign for their predators which provide protection for them to survive (Hickman et al., 2018).

Based on the IUCN Red List of Threatened Species (2021), all of the species recorded were listed as the Least Concern (LC) species. Least Concern (LC) species is considered as not being a focus of species conservation by the International Union for Conservation of Nature (IUCN). However, Pasir Gudang City Council should monitor the anthropogenic activities especially in manufacturing and industrial activities that occur around Pasir Gudang city such as by ensuring these activities are restricted to the environmental friendly aspect. Findings from this study can ensure that the authorities are always alert and aware towards the existence of the diversity and variety of Lepidoptera present in that area so that they can protect and contribute to a better conservation and management system.

As a conclusion, a total of nine species from three families of order Lepidoptera were successfully recorded from three different urban parks at Pasir Gudang, Johor. Specifically, there were seven species of Lepidoptera recorded at Laman Rekreasi Sungai Buluh, four species were recorded at Tasik Dahlia and six species were recorded at Taman Bandar. The most abundant species at Laman Rekreasi Sungai Buluh was *Junonia hedonia* with 13 individuals and followed by *Appias libythea* with seven individuals. The most abundant species recorded at Tasik Dahlia was *Zizula hylax* with 16 individuals. The most abundant species recorded at Taman Bandar were *Elymnias hypermnestra* and *Eurema sari* with 11 individuals each. Diversity of Lepidoptera at Laman Rekreasi Sungai Buluh showed the highest compared to the Tasik Dahlia and Taman Bandar. This were supported by the number of abundance species recorded, number of individuals recorded, statistical analysis (t-test) and diversity indices. There was a significant different in Lepidoptera diversity between the study sites. The sampling efforts at three study sites were enough and no additional species would be documented even with the additional of sampling sessions for all study sites.

The findings from this study may aid the conservation efforts to protect Lepidoptera species in many urban parks in Pasir Gudang city. To increase the population of Lepidoptera around the urban parks area, the management of the urban parks can do modification in types of vegetation especially by adding more native flowering plants such as *Clerodendrum paniculatum, Begonia obliqua* and *Bougainvillea spectabilis* because these plants can be one of the main food source for Lepidoptera. The finding from this study may also be a baseline information to the future study by other researchers. The results from this study may be used to increase the public awareness and knowledge about how important to manage anthropogenic activities and to manage the landscape of the urban parks to ensure the vegetations that are planted in the urban parks are friendly to Lepidoptera community and indirectly will preserve and conserve the Lepidoptera species in Johor.

## ACKNOWLEDGEMENTS

A highly gratitude to the Pasir Gudang City Council, Johor for giving the permission to carried out this research. We also would like to thank Mr. Muhammad Azlan Syah Bin A Aziz who had assisted us during the field work.

## Notes

### Competing Interest Statement

The authors have declared no competing interest.

